# OmicsSankey: Crossing Reduction of Sankey Diagram on Omics Data

**DOI:** 10.1101/2025.06.13.659656

**Authors:** Shiying Li, Bowen Tan, Si Ouyang, Zhao Ling, Miaozhe Huo, Tongfei Shen, Jingwan Wang, Xikang Feng

## Abstract

In bioinformatics, Sankey diagrams have been widely used to elucidate complex biological insights by visualizing gene expression patterns, microbial community dynamics, and cellular interactions. However, computational scalability remains a challenge for large-scale biological networks. In this work, we present OmicsSankey, a novel formulation of the layout optimization problem for Sankey Diagrams that employs eigen decomposition as a closed-form solution, addressing graph disconnection through a teleportation mechanism that enhances connectivity and stabilizes eigenvector solutions. Experimental results on synthetic datasets with varying layers and nodes validate the efficacy of OmicsSankey compared to state-of-the-art layout-optimizers. Improving the Sankey layouts for Cell Layers, BioSankey, and Sequence Flow further validates OmicsSankey in enhancing the interpretability of biological insights.

## 1. Introduction

Sankey diagrams are powerful visualization tools for representing flows among entities in various systems, where the width of each flow corresponds to its magnitude. In bioinformatics, these diagrams have become increasingly relevant, particularly in multi-omics applications such as single-cell RNA sequencing (scRNA-seq) analysis, metagenomics, and gene expression studies [1, 2, 3]. Tools like Cell Layers [4], BioSankey [5], and Sequence Flow [6] leverage Sankey diagrams to elucidate complex biological insights by visualizing gene expression patterns, microbial community dynamics, and cellular interactions. By examining these data flows, researchers can interpret biological relationships, identify key regulatory mechanisms, and derive meaningful conclusions from high-dimensional datasets.

Layout optimization is crucial for accurately representing complex biological relationships [7]. As the complexity of biological data increases—reflected in the growing number of vertices and edges—the challenge of achieving optimal layouts becomes even more significant. Tools like Cell Layers, BioSankey, and Sequence Flow rely on clear visualizations to extract meaningful insights. The main tasks involved in enhancing a Sankey layout include minimizing crossings, shortening edge lengths, and maintaining aesthetic clarity [8]. Excessive edge crossings can obscure relationships, complicating the interpretation of biological patterns and potentially leading to misinterpretations. Thus, reducing edge crossings is vital for enhancing interpretability, enabling researchers to draw accurate conclusions from complex datasets.

The NP-hard nature of crossing minimization [9] has spurred the development of heuristic solutions. Barycentric approaches dominate current methodologies, including Sugiyama’s foundational BC method [10], which positions vertices at the averages of adjacent nodes, and its median-based variant that offers 3-approximation guarantees [11]. Although linear programming refinements [8] and integer linear programming (ILP) formulations [12] address weighted edge scenarios, computational scalability remains a challenge for large-scale biological networks. Concurrently, spectral drawing techniques leveraging Laplacian eigenvectors [13, 14] show untapped potential for crossing reduction despite their proven efficacy in graph embedding. In this work, we present OmicsSankey, a novel formulation of the lay-out optimization problem for Sankey diagrams that integrates spectral techniques with barycentric ordering. Our approach derives eigen decomposition as a closed-form solution while addressing potential graph disconnection through a teleportation mechanism that enhances connectivity and stabilizes the resulting eigenvector solutions. Additionally, OmicsSankey incorporates a refinement step that allows for adaptive vertex placement within defined ranges to further minimize edge crossings. This comprehensive method combines advanced graph-theoretic principles with geometric insights, providing an effective solution for visualizing complex data flows in bioinformatics. We evaluate OmicsSankey on synthetic datasets with varying layers and nodes, demonstrating its competitive performance and runtime relative to the optimal ILP method. Extensive comparisons against the ILP method, the barycentric method, and a state-of-the-art heuristic method on real datasets further validate the efficacy of OmicsSankey in enhancing the interpretability of biological insights.

## 2. Methods and materials

### 2.1. Problem Definition

In a Sankey diagram, entities are organized into vertical layers, with links of varying widths representing quantities connecting adjacent entities. To accommodate links between non-successive layers, dummy entities may be introduced. This results in a *k*-layered graph *G*, where each entity corresponds to a vertex *u*. Let *V*_*i*_ denote the set of vertices in the *i*-th layer, and _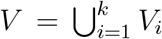_ be the overall vertex set. An undirected edge ⟨*u, v*⟩ exists if a link connects *u* and *v*, with weight *w*_⟨*u*,*v*⟩_ equal to the link’s width. The edge set is denoted as *E*, with *E*_*i*_ representing edges between the *i*-th and (*i* + 1)-th layers.

For example, in the first layer of Figure 1A, entities A, B, and C correspond to vertices *a, b, c*. A link from E to X with quantity *Q* = 7 gives *w*_⟨*e*,*x*⟩_ = 7, while a link from F to Y with *Q* = 9 results in *w*_⟨*f*,*y*⟩_ = 9. A long link from C to Y requires a dummy vertex 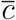 in layer 2, represented by edges 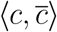 and 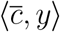. Dashed lines indicate virtual vertical boundaries for the layers.

**Figure 1.**
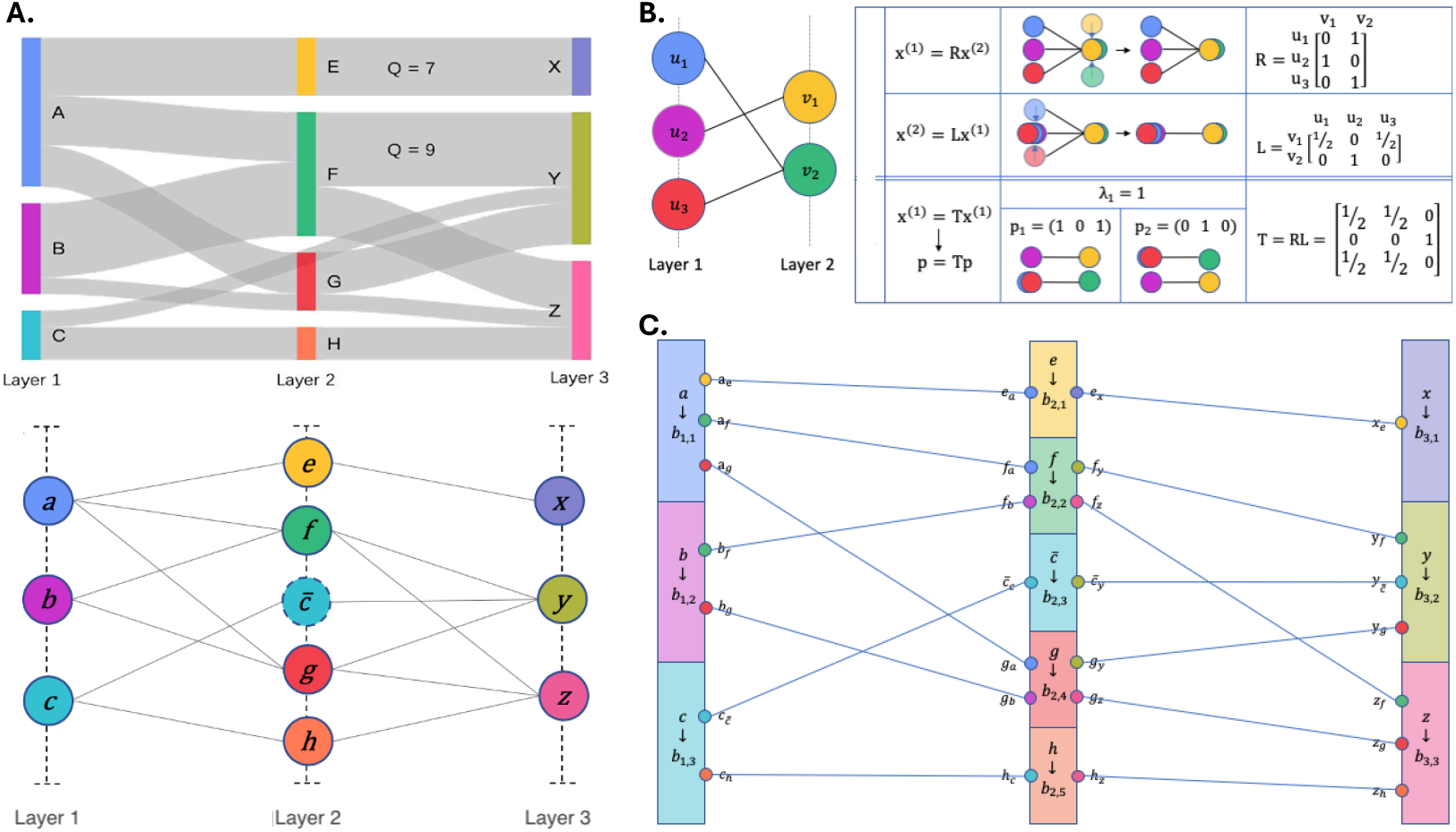
OmicsSankey workflow. **A**. A 3-layer Sankey diagram (top) and the corresponding 3-layer graph *G* (bottom). **B**. Stage one workflow. A case demonstrating the calculation of barycentre ordering. Left: graph representation of a two-layer Sankey diagram with all edge weights equal to 1. Upper right: iterative steps of updating vertex positions to obtain a barycentre ordering and resultant graphs. Upper left: the Sankey characteristics matrix and the graphs corresponding to the two Eigenvectors of *λ*_1_ = 1. **C**. Stage two workflow. The micro view of *G* in (A) with the original ordering. Based on the second layer ordering, vertex *f* has the second-highest rank and is assigned to the second block *b*_2,2_. In Micro-Dissection, the two vertices *a* and *b* in *N*_*L*_(*f* ) are connected to points *f*_*a*_ and *f*_*b*_ in block *b*_2,2_. Notably, *a* having a higher rank than *b* in the ordering leads to *p*(*f*_*a*_) *> p*(*f*_*b*_).

Vertices in each layer are linearly ordered along a virtual vertical line, with each vertex *v* ∈ *V*_*i*_ having a coordinate *p*(*v*). The ordering, denoted *σ*_*i*_, ranks vertices based on *p*(*u*) in descending order, resulting in *σ*(*G*) = {*σ*_1_, … , *σ*_*k*_ }.

A crossing occurs between edges ⟨*u, v*⟩ and ⟨*u*^′^, *v*^′^⟩ if *σ*_*i*_(*u*) *> σ*_*i*_(*u*^′^) and *σ*_*i* +1_(*v*) *< σ*_*i*+1_(*v*^′^) or vice versa. The weight of a crossing is the product of the weights of the crossed edges. Our goal is to find a layout *σ*(*G*) that minimizes weighted crossings, represented as *W* (*σ*(*G*)), while non-weighted crossings are denoted as *K*(*σ*(*G*)). The computation of *W* (*σ*(*G*)) follows a modified approach based on the method for *K*(*σ*(*G*)) from [15]. The weighted crossings of *E*_*i*_ with respect to *σ*_*i*_ and *σ*_*i*+1_ are calculated as:

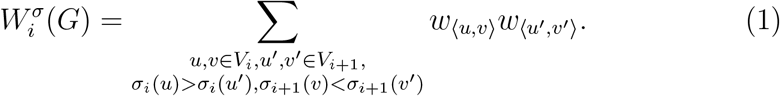

Thus, the total weighted crossing number for *G* is 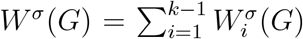, and we aim to determine an ordering that minimizes the total weighted crossings.

We define the ordering of a barycentric l ayout a s *barycentric ordering*, where each vertex’s position is determined by the weighted average of its adjacent vertices. For a vertex *u*, let *N* (*u*) be its neighbors. For *u* ∈ *V*_*i*_ with *i* ∈ [2, *k* − 1], *N* (*u*) is partitioned into left and right sets *N*_*L*_(*u*) and *N*_*R*_(*u*) from the (*i* − 1)-th and (*i* + 1)-th layers, respectively. For vertices in the first or last layer, *N* (*u*) includes only one-sided neighbors.

The barycentre of the left neighboring set for vertex *u* is given by:

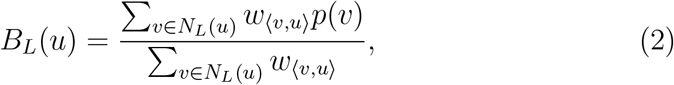

if *i* ∈ [2, *k*], and *B*_*L*_(*u*) = *B*_*R*_(*u*) if *i* = 1. Similarly, the barycentre of *B*_*R*_(*u*) is defined by replacing *N*_*L*_(*u*) with *N*_*R*_(*u*) if *i* ∈ [1, *k* − 1], and *B*_*R*_(*u*) = *B*_*L*_(*u*) if *i* = *k*. The two-sided barycentre of *u* is then computed as the average of the two one-sided barycentres. Consequently, finding a barycentric ordering involves determining the positions of vertices based on the barycentres of their adjacent vertices.

### 2.2. OmicsSankey on the Sankey diagrams

#### 2.2.1. Formulating barycentric layouts with a transition matrix

We define the *position vector* **x**_*i*_ of the *i*-th layer as *x*_*i*,*u*_ = *p*(*u*) for all *u* ∈ *V*_*i*_. For *i* ∈ [1, *k* − 1], the position vector satisfies *x*_*i*+1,*v*_ = *B*_*L*_(*v*) for each *v* ∈ *V*_*i*+1_, or **x**_*i*+1_ = **L**_*i*_**x**_*i*_, where **L**_*i*_ is the *i*-th *left transition matrix* derived from the weighted adjacency matrix **M**_*i*_ between *V*_*i*_ and *V*_*i*+1_. For *i* ∈ [2, *k*], we express **x**_*i*−1_ as *x*_*i*−1,*u*_ = *B*_*R*_(*u*) for *u* ∈ *V*_*i*−1_, with the *right transition matrix* **R**_*i*_ also derived from **M**_*i*_. Thus, **x**_*i*−1_ = **R**_*i*_**x**_*i*_.

By propagating a given position vector **x**_1_ through the product of the modified left transition matrices from **L**_*k*−1_ to **L**_1_, we position all vertices in all but the first layers at their left barycentres based on **x**_1_. This process is equivalent to the equation **x**_*k*_ = (**L**_*k*−1_**L**_*k*−2_ … **L**_1_)**x**_1_. Conversely, given the position vector **x**_*k*_, we reverse the process using **x**_1_ = (**R**_1_**R**_2_ … **R**_*k*−1_)**x**_*k*_, thereby locating vertices in all but the last layer at their right barycentres based on **x**_1_. Consequently, we calculate **x**_*k*_ from an initial position vector **x**_1_, which we then update reversely to obtain **x**_1_. This iterative process continues until convergence, meaning that both **x**_1_ and **x**_*k*_ remain unchanged in an iteration. Specifically, this results in each vertex *u* ∈ *V* locating at both its left and right barycentres, i.e., *B*(*u*) = *B*_*L*_(*u*) = *B*_*R*_(*u*). We achieve a barycentric ordering through this iterative back-and-forth process.

Let **L** = **L**_*k*−1_**L**_*k*−2_ … **L**_1_ of size |*V*_*k*_| × |*V*_1_| and **R** = **R**_1_**R**_2_ … **R**_*k*−1_ of size |*V*_1_| × |*V*_*k*_| . We simplify this process as **x**_1_ = **RLx**_1_ := **Tx**_1_. Here, we define the *Sankey characteristics matrix* **T** = **RL**. By definition, both **L** and **R** are stochastic matrices, meaning they are non-negative real square matrices with each row summing to 1. Consequently, their product **T** is also a stochastic matrix.

The solution to the equation **p** = **Tp** via eigendecomposition of matrix **T** yields the Eigenvector **p**_1_ corresponding to the Eigenvalue *λ*_1_ = 1, which is invariant under **T**. Setting **x**_1_ = **p**_1_, we can derive all other position vectors using **x**_*i*+1_ = **L**_*i*_**x**_*i*_ and ultimately obtain the ordering *σ*(*G*).

#### 2.2.2. Eigen-decomposition on enhanced adjacency matrix with teleport operations

A problem with the above formulation is that Sankey diagrams may be ill-defined or disconnected, leading to a Sankey characteristics matrix **T** that is not strongly connected. According to the Fundamental Theorem of Markov chains, the Eigenvector **p**_1_ exists but is non-unique, making it ambiguously defined. Figure 1B illustrates the iterative steps and l ayouts d erived from the Eigenvectors corresponding to the Eigenvalue *λ*_1_ = 1.

To address the disconnectedness of graph *G*, we introduce a teleport operation by adding a random matrix 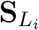 with normalized rows into the equation **x**_*i*+1_ = **L**_*i*_**x**_*i*_, scaled by a small factor *α* ∈ [0, 1]. This results in the modified equation 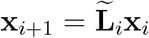, where 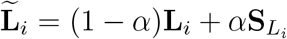 . A similar approach for the right transition matrix **R** connects the graph *G*, refining 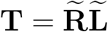 while preserving stochastic properties.

For a connected graph, Koren showed that barycentric ordering can only be achieved through a degenerate solution that places all vertices at the same location [14]. Conversely, the second-largest sign-less eigenvalue *λ*_2_ provides an eigenvector **p**_2_ as a heuristic solution to **p** = **Tp**. The sign of *λ*_2_ is irrelevant, as it only inverts the ordering without affecting edge crossings. Liu et al. applied this concept to the NP-complete state clustering problem [16], where the eigenvector corresponding to *λ*_2_ offers a satisfactory approximation for clustering. Similarly, in our framework, **p**_2_ serves as a competent alternative to **p**_1_.

The teleport operation introduces randomness into **T**, affecting the final ordering. To mitigate variability, we implement an *N-selection* process, repeating the procedure *N* times and recording the output ordering *σ*_*t*_(*G*) and _9_ its weighted crossings *W* (*σ*_*t*_(*G*)). The ordering with the minimum weighted crossings is selected as the final result, denoted as 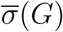. This method benefits from efficient eigen decomposition techniques for computing *λ*_2_ and its corresponding eigenvector, with the dimension of **T** being |*V*_1_|. Although teleportation introduces randomness, a larger *N* typically reduces the weighted crossing number 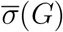. We designate 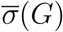 as *Eigen-centric ordering*, the near-barycentre positions as *Eigen-centres*, and the entire process as *Eigen-Production*.

#### 2.2.3. Refining vertex placement with Micro-Dissection

While Eigen-centric ordering provides a satisfactory alternative to barycentric ordering, it can be refined through a technique termed *Micro-Dissection*. Instead of representing each vertex as a single point, we allow each vertex to assign distinct position values within a specified range to its adjacent vertices for barycentre calculations. Each vertex is allocated a non-overlapping range, referred to as a *block*, within its layer. A microview is achieved by using a point within this range to represent the endpoint of each edge connected to the vertex. For any *v* ∈ *N* (*u*), we denote the point on edge ⟨*u, v*⟩ as *u*_*v*_ and its position as *p*(*u*_*v*_).

Given an ordering *σ*(*G*), the position *u*_*v*_ depends on the ranks and positions of *u* and *v* within their respective layers. We divide the *i*-th layer into |*V*_*i*_| blocks of equal height, assigning vertex *u* with rank *σ*_*k*_(*u*) to the *σ*_*k*_(*u*)-th block from top to bottom. Thus, the rank of *u* determines the range of *u*_*v*_. The value of *u*_*v*_ within the block is determined by the position of *v*, specifically *p*(*v*_*u*_). The *λ*-th block in the *i*-th layer has a base *b*_*i*,*λ*_ = (|*V*_*i*_| − *λ*)*/*|*V*_*i*_| and height *h*_*i*_ = 1*/*|*V*_*i*_|. The initial position of *u*_*v*_ is derived from *u*’s rank in the Eigen-centric ordering *σ*(*G*). For vertices *v* and *v*^′^ ∈ *N*_*L*_(*u*), the ordering of points *u*_*v*_ and *u*_*v ′*_ aligns with that of *v* and *v*^′^. The points for vertices in *N*_*L*_(*u*) are distributed evenly within the block, dividing it into |*N*_*L*_(*u*)| + 1 equal segments:

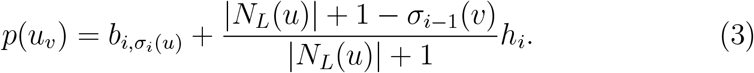

Points corresponding to vertices in *N*_*R*_(*u*) follow a similar process, substituting |*N*_*R*_(*u*) | for |*N*_*L*_(*u*)|. Notably, points for vertices in different neighboring sets may share a position.

Using the Eigen-centric ordering 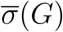, we assign blocks and points for all vertices based on their ranks within the layer. We then calculate the initial positions for all points following the aforementioned equation. The iteration begins by computing the barycentres of vertices in the middle layers, from *i* = 2 to *i* = *k* − 1. For example, *B*_*L*_(*u*) is defined by replacing *p* (*v*) in Equation 2 with *p*(*v*_*u*_).

The two-sided barycentre of *u* is the average of the one-sided barycentres. The calculated barycentres for vertices in the second layer yield a new ordering *σ*_2_, arranged in descending order of barycentres. Based on the new *σ*_2_, we reassign blocks for each vertex. The new position of point *u*_*v*_ is influenced by the position of *v*_*u*_ given 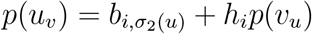.

The calculations for the remaining middle layers proceed similarly. For each vertex in the last layer, its barycentre equals its left barycentre. After updating the ordering, blocks, and points for the last layer, we iterate backwards, repeating the operations on the middle layers from *i* = *k* − 1 to *i* = 2. The procedure for the first layer mirrors that of the last layer, completing one iteration.

After each iteration, we compute and record the weighted crossings of the resulting orderings. Following *M* predefined iterations, we select the ordering with the minimum weighted crossings as the final result of the Micro-Dissection process. This approach is justified, as while this process converges towards a barycentric ordering and consequently a barycentric layout, a barycentric layout is not necessarily optimal. Thus, it is feasible to discover a layout with fewer weighted crossings than that of the resulting barycentric layout after *M* iterations.

### 2.3. Simulation construction

The process of generating simulation datasets involves systematically constructing Sankey diagrams with varying numbers of nodes and links. The scale of each simulated dataset is controlled by the number of layers *n* and the average number of vertices per layer 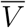. Following the dataset scale, we distribute nodes into distinct layers, each with a predetermined capacity, allowing us to simulate resource flow between nodes while adhering to capacity constraints. Edges are constructed using a probabilistic mechanism that dynamically updates the nodes’ remaining capacities as connections are formed. For each unique pair of *n* and 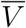, we generated ten random test cases, with the total number of edges serving as an estimate of complexity after introducing the random effect. The implementation of the ILP method utilizes the PuLP library in [17].

### 2.4. Evaluation metrics

In real datasets, the weighted and non-weighted crossings serve as performance measurements. For simulated cases, we evaluate the effectiveness of our modified method by comparing the resultant weighted crossing numbers to the theoretical optimum using a Δ metric. To account for the varying complexities of the datasets, we standardized the weighted crossing numbers relative to the worst-case scenario, where every two edges in *E*_*i*_ cross for any *i* ∈ [1, *n*]. Specifically, let *W*_*opt*_(*G*) represent the weighted crossings from the optimal layout and *W* (*G*) from the worst layout. We define 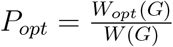as the standardized optimum. In our cases, since *W*_*opt*_(*G*) = 0, we have *P*_*opt*_ = 0. The weighted crossing number from OmicsSankey is denoted as *W* ^′^(*G*), and we define 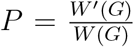 as our standardized measure. The difference from the optimal layout is captured by Δ = *P* − *P*_*opt*_.

## 3. Result

### 3.1. Benchmarking with state-of-the-art methods on synthetic datasets of different scales and real datasets

We conducted a series of tests to evaluate the performance and running time of OmicsSankey across varying complexity levels, defined by the number of layers (*n*) and the average number of vertices per layer 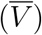. For each unique pair of *n* and 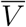, we generated ten random test cases, with the total number of edges serving as an estimate of complexity. All test cases were executed with *α* = 0.01, *N* = 100 for both Eigen-Production and Micro-Dissection. We recorded the weighted crossings for both the Eigen-centric and final ordering, alongside the optimal results from the ILP method for comparative analysis. Performance was measured using Δ from the previous subsection, and we also documented the running times for both the proposed and ILP methods

The results, summarized in Figure 2, indicate that all Δ values from the Eigen-centric ordering remain below 0.05, demonstrating satisfactory performance. Following Micro-Dissection, the mean Δ values across all datasets for any pair of *n* and *V* drop below 0.02, confirming the effectiveness of OmicsSankey across varying complexities.

**Figure 2.**
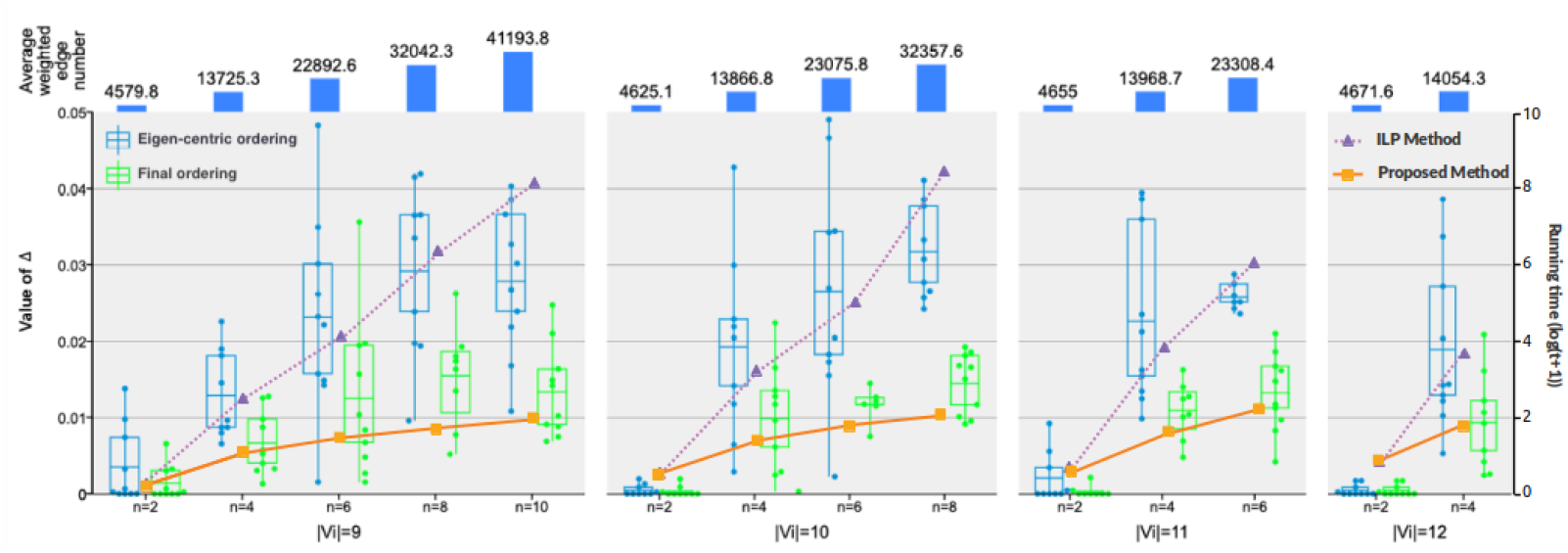
Results of performance and running time in second. The Δ values are shown by the box plots with the left axis, and the running time is shown by the lines with the right axis.

Regarding computational efficiency, OmicsSankey exhibits comparable performance to the ILP method on small datasets (with two layers). However, on larger datasets (with more than two layers), OmicsSankey significantly outperforms the ILP method in terms of speed, highlighting its suitability for practical applications in complex scenarios.

Subsequently, we compare OmicsSankey with the state-of-the-art method on real world dataset, starting with the heuristic method by Alemasoom *et al*. [8] using Canada’s 1978 energy usage dataset. For Eigen-centric ordering, we set *α* = 0.1 and *N* = 100, while for Micro-Dissection, we maintained *α* = 0.1 and set *M* = 50. Both orderings were evaluated using weighted and non-weighted crossing measurements, with visual representations in Sankey diagrams.

The results indicate that OmicsSankey outperformed the heuristic method, achieving lower scores in both weighted (82.79) and non-weighted crossings (157) compared to the heuristic’s scores of 146.77 and 279 (Figure 3, upper row). Notably, edge crossings were more pronounced in the last two layers of the heuristic ordering, whereas our method effectively mitigated these crossings.

**Figure 3.**
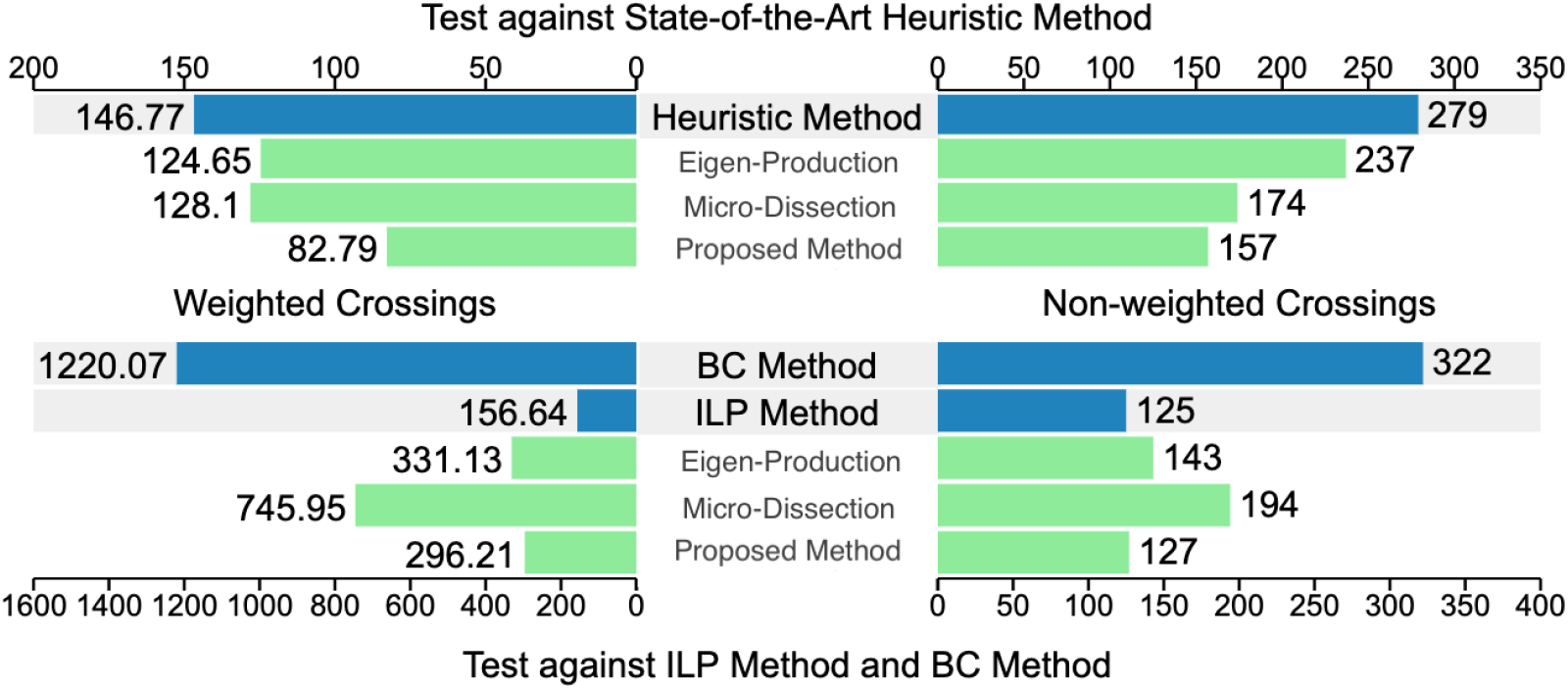
Two back-to-back bar charts showing the weighted (left) and non-weighted (right) crossings of the output layouts in 1978 Canada’s energy usage dataset (top) and the world greenhouse gas emissions dataset (bottom).

In individual stages, the Micro-Dissection stage showed slightly lesser performance, with Eigen-centric ordering yielding lower weighted crossings (124.65) and higher non-weighted crossings (237) compared to Micro-Dissection’s scores of 128.1 and 174. Micro-Dissection demonstrated a nearly 30% reduction in both crossing measurements, achieving the fewest weighted crossings at the 35th iteration.

We also compared our method with BC and ILP methods using World Greenhouse Gas Emissions data from the World Resources Institute [18]. For this comparison, we set *α* = 0.01 and *N* = 100 for Eigen-Production, and *α* = 0.1 and *M* = 50 for Micro-Dissection. The results, summarized in Figure 3, show that OmicsSankey outperformed the BC method, recording significantly lower weighted (156.64) and non-weighted crossings (125) compared to BC’s scores of 1220.07 and 322. The Eigen-centric ordering also surpassed the BC method with scores of 331.13 and 143.

Additionally, OmicsSankey’s final ordering closely approximated the optimal layout from the ILP method, introducing only two additional intersections and resulting in a slight increase in weighted crossings (139.57). While Micro-Dissection alone outperformed the BC method, it showed a larger discrepancy from the ILP method and Eigen-centric ordering, with crossings of 745.95 and 194, respectively. Nonetheless, applying Micro-Dissection to Eigen-centric ordering significantly enhanced performance, demonstrating its efficacy in reducing crossings. Overall, these results affirm that Micro-Dissection effectively improves upon the initial Eigen-centric ordering.

### 3.2. OmicsSankey generating optimal Sankey layouts that reveal node relation dynamics

In bioinformatics analyses, Sankey diagram is often used for represent hierarchy-like data, where links represent membership changes along an ordered condition. For example, Cell Layers visualizes the flow of cells between different cluster assignments across multiple resolutions of analysis. Each layer represents a specific resolution used for clustering, with nodes depicting distinct cell clusters. The node sizes scale with cluster sizes, and the edge widths encode the magnitude of cell transitions between clusters. Similarly, BioSankey illustrates the taxonomic distribution of the most abundant genera from the tongue tissue [19]. The nodes represent specific taxa (*e*.*g*., phyla or genera), while the columns group these nodes by taxonomic rank from higher levels on the left to lower levels on the right. The edge crossing hinders the identification of taxonomic distribution along taxonomic ranks.

The original Sankey layouts from Cell Layers and BioSankey can suffer from substantial edge crossings, making it difficult to trace cell membership across resolutions, or taxa relation in specification. OmicsSankey obtained optimal layouts with zero crossing for Cell Layers (Figure 4A-B), with five crossing and 20,397 weighted edge crossing, and BioSankey, with four edge crossings and 67,789,803 weighted edge crossing (Figure 4C-D).

**Figure 4.**
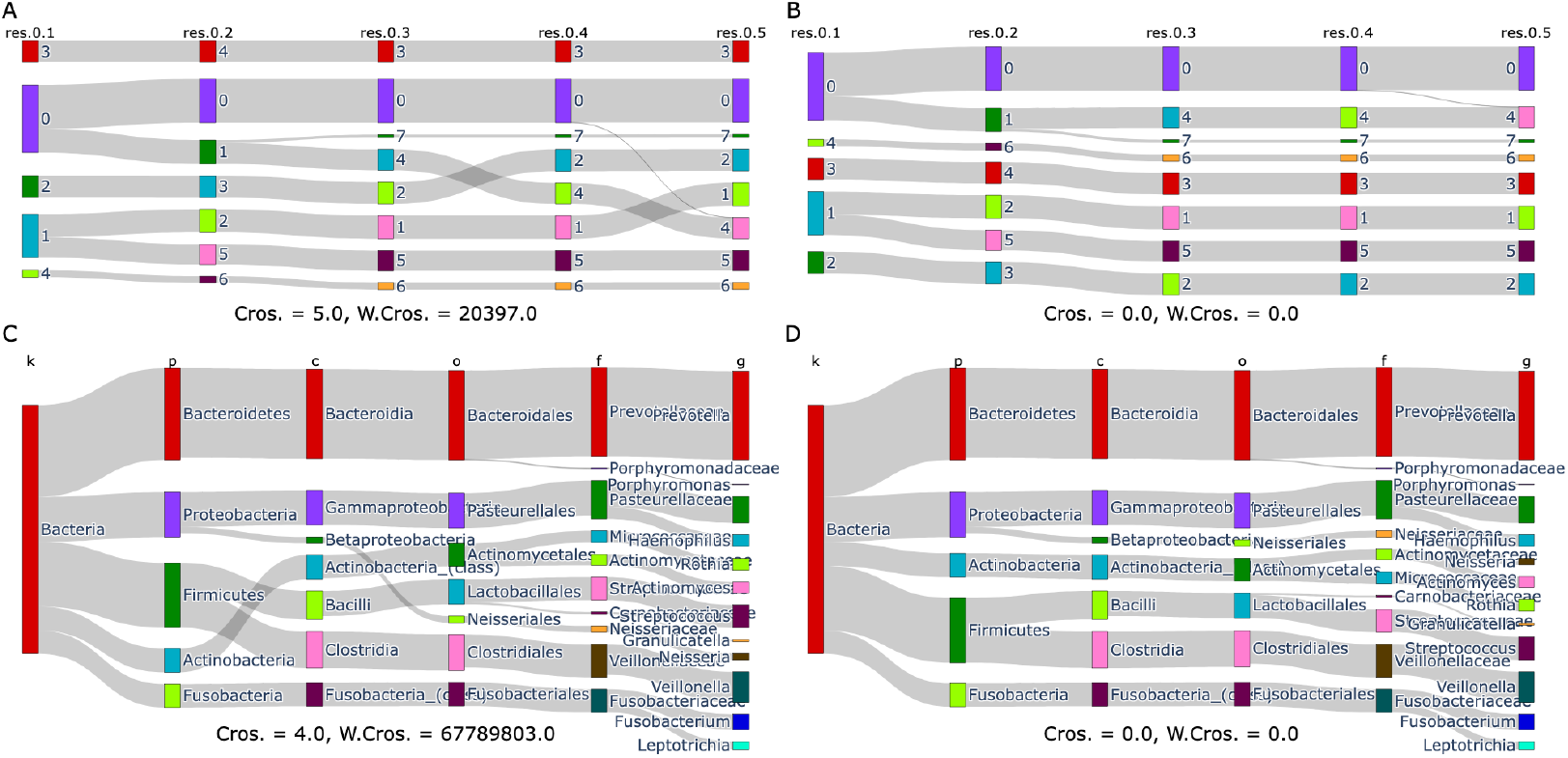
Comparison of original (left) and optimized (right) Sankey diagram layouts. **A-B**. Cell Layers PBMC multi-resolution analysis. **C-D**. BioSankey taxonomic distribution of the most abundant genera from the tongue tissue.

This optimized layout provides several key advantages. Firstly, the removing of edge cross allows for identifying nodes that are invariant against the condition. For Cell Layers, such nodes represent conserved clusters across resolutions, which could indicate real biological structures or cell types rather than noise or over-clustering. For BioSankey, the conserveness of actinobacteria becomes apparent when OmicsSankey resolved all crossing.

The OmicsSankey-optimized layouts also allows for clear observation of how node relation changes across various conditions. The optimized Cell Layers clearly depicts how cell clusters partition and change across resolutions. For example, we can now clearly see how cluster 0 from resolution 0.1 further partitions into clusters 0 and 1 in resolution 0.2, with the latter then splitting into clusters 7 and 4 in resolution 0.3. This enhanced visibility of cluster membership changes facilitates a deeper understanding of the underlying cell population dynamics, or cell hierarchies [20]. The taxonomic Sankey diagram also benifits from the optimized layout, with clearer specification ralation from fusobacteria to bacilli and clostridia.

Moreover, the optimized layouts better represent the similarity between nodes within the same layer. By minimizing edge crossings, nodes that are more closely related, such as clusters 4 and 0 in resolution 0.5 in the Cell Layers graph, are positioned in closer proximity. This spatial arrangement provides additional visual cues about the relative similarity between cell clusters, complementing the information conveyed by the edge connections. Similarly, the proximity of pasteurellales and neisseriales after OmicsSankey optimization also reflects their resemblance.

In summary, the OmicsSankey optimization of the Sankey diagram layouts significantly improves their interpretability. The clear depiction of cluster partitioning and taxonomic specification empower researchers to gain deeper insights into the complex dynamics of the studied populations.

### 3.3. OmicsSankey reveals novel insights for partial order alignment Sankey diagram

Partial Order Alignment (POA) has been put forward as an alternative method for multiple sequence alignment (MSA) [21]. In a POA Sankey diagram, nodes are residue clusters of the aligned sequences, and links connect successive residues in these sequences. In protein POA Sankey diagrams, the limited number of common nodes of related sequences leads to a larger number of nodes per layer. Such complexity in the POA graph layout can cause difficulty in recognizing and assessing similarity between sequences. While Sequence Flow attempted to address this issue by combining amino acids with similar physicochemical properties [6], OmicsSankey presents as an alternative solution for reducing layout complexity.

Applying OmicsSankey to the POA graph for the msl ref7 dataset from the BAliBASE 2 [22] reduced crossing from 91 to 60, and weighted crossing from 182 to 80 (Figure 5). The improved layout presents a clearer view on sequence similarity while maintained amino acid information. For example, Isoleucine (I) is most similar to Proline (P) and Alanine (A) due to shared hydrophobic and non-polar characteristics, and is likely to be grouped as one node in Sequence Flow. However, in the fifth layer, the difference in the left and right neighbors of the three nodes can reveals subtle variations in sequences. Such insights, while possibly lost by simplifying graph nodes as proposed by Sequence Flow, can be retained by optimizing the Sankey layout with OmicsSankey.

**Figure 5.**
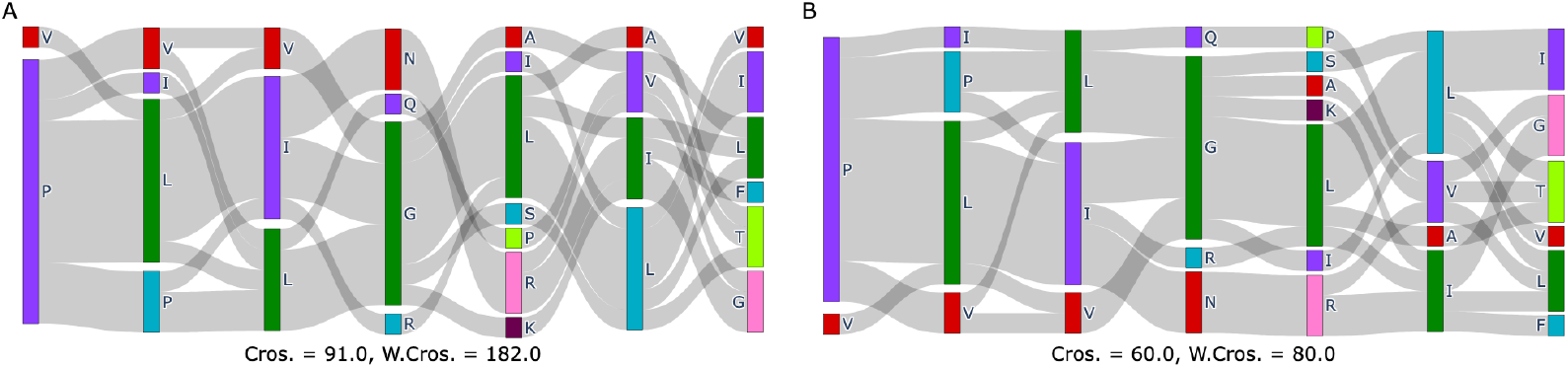
Comparison of Sequence Flow (**A**) and optimized (**B**) Sankey diagram layouts for partial order alignment on msl ref7 alignment.

Furthermore, the OmicsSankey-optimized layout can provide novel insight in to sequence variations with the edge crossings between consecutive layers after optimization. In the original POA Sankey siagram, the number of crossing between consecutive layers does not always convey biological information, as they can be the results of a non-optimal layout (such as the fourth and fifth layers). The OmicsSankey layout, on the other hand, can truthfully represent biological relevant information towards the level of sequence variations.

## 4. Conclusion and Further Work

We reformulate the BC method as the Eigendecomposition of a Sankey characteristics matrix with a connectivity problem. We refine the Sankey characteristics matrix with a teleport operation and obtain its second-largest sign-less Eigenvalue through Eigendecomposition. While the final ordering is already better than those of existing heuristic methods, further improvements can be made to the Sankey characteristics matrix, such as calculating the median instead of the average for barycentres.

The results from our analyses demonstrate that OmicsSankey effectively enhances the visualization of complex data through optimized Sankey diagram layouts. By outperforming existing methods in both performance and computational efficiency, particularly on larger datasets, OmicsSankey proves to be a valuable tool for handling intricate scenarios. Its ability to reduce edge crossings significantly improves the interpretability of bioinformatics data, allowing for clearer identification of conserved clusters and taxonomic distributions. Additionally, in the context of POA, OmicsSankey provide novel insights by enhancing the clarity of sequence similarity while preserving important amino acid information. These advancements facilitate a deeper understanding of underlying dynamics in biological data, showcasing OmicsSankey’s potential for broad applications in bioinformatics.

We acknowledge several limitations in OmicsSankey that challenge optimal layouts for Sankey diagrams. First, disconnected graphs can introduce variability due to randomness from the teleport operation. Second, reliance on eigenvalues, particularly the second-largest eigenvalue, may lead to sub-optimal clustering in complex graph structures. To address these issues, we developed the Micro-Dissection technique, allowing each vertex to occupy a non-overlapping range within its layer. This granularity enables finer adjustments based on local neighborhood structures, promoting stability and consistency across layouts. Additionally, by combining barycentric calculations with real-time adjustments based on local edge weights, we enhance connectivity and improve visualization quality by minimizing edge crossings in a context-sensitive manner.

## Funding statement

This study was funded by the National Natural Science Foundation of

China (32300527), and the Basic Research Programs of Taicang, 2024 (TC2024JC43).

## CRediT authorship contribution statement

**Xikang Feng:** Writing – review & editing, Supervision, Resources, Conceptualization, Project administration, Methodology, Investigation, Funding acquisition. **Shiying LI:** Writing – original draft, Visualization, Validation, Software, Formal analysis, Data curation. **Bowen TAN:** Writing – review & editing, Methodology. **Si Ouyang:** Writing – review & editing, Software. **Zhao Ling:** Writing – review & editing. **Miaozhe Huo:** Writing – review & editing. **Tongfei Shen:** Writing – review & editing. **Jingwan Wang:** Writing – review & editing.

## Code availability

The source code and analysis scripts are freely available in https://github.com/deepomicslab/OmicsSankey.

## Acknowledgments

We thank Dr. Yen Kaow Ng for his constructive comments that greatly improved this manuscript.

## References

[1] H. Liu, Z. Deng, B. Yu, H. Liu, Z. Yang, A. Zeng, M. Fu, Identification of slc3a2 as a potential therapeutic target of osteoarthritis involved in ferroptosis by integrating bioinformatics, clinical factors and experiments, Cells 11 (21) (2022) 3430.

[2] M. Zhang, T. Ge, Y. Zhang, X. La, Identification of mark2, ccdc71, gata2, and klrc3 as candidate diagnostic genes and potential therapeutic targets for repeated implantation failure with antiphospholipid syndrome by integrated bioinformatics analysis and machine learning, Frontiers in Immunology 14 (2023) 1126103.

[3] Y. Xing, Y. Wang, M. Pi, R. Zhang, J. Yang, T. Wen, Syndrome evolution and chinese herb formula regularity of tcm heat syndrome in type 2 diabetes mellitus: complex network community discovery algorithm and sankey diagram visualization, in: 2020 IEEE International Conference on Bioinformatics and Biomedicine (BIBM), IEEE, 2020, pp. 1617–1621.

[4] A. P. Blair, R. K. Hu, E. N. Farah, N. C. Chi, K. S. Pollard, P. F. Przytycki, I. S. Kathiriya, B. G. Bruneau, Cell layers: uncovering clustering structure in unsupervised single-cell transcriptomic analysis, Bioinformatics Advances 2 (1) (2022) vbac051.

[5] A. Platzer, J. Polzin, K. Rembart, P. P. Han, D. Rauer, T. Nussbaumer, Biosankey: visualization of microbial communities over time, Journal of Integrative Bioinformatics 15 (4) (2018) 20170063.

[6] K. Zdab-lasz, A. Lisiecka, N. Dojer, Sequence flow: interactive web application for visualizing partial order alignments, BMC genomics 25 (1) (2024) 973.

[7] F. Paduano, A. G. Forbes, Extended linesets: a visualization technique for the interactive inspection of biological pathways, in: BMC proceedings, Vol. 9, Springer, 2015, pp. 1–13.

[8] H. Alemasoom, F. Samavati, J. Brosz, D. Layzell, Energyviz: an interactive system for visualization of energy systems, The Visual Computer 32 (3) (2016) 403–413. doi:10.1007/s00371-015-1186-8. URL https://doi.org/10.1007/s00371-015-1186-8

[9] M. R. Garey, D. S. Johnson, Crossing number is NP-complete, SIAM Journal on Algebraic Discrete Methods 4 (3) (1983) 312–316. doi:10.1137/0604033.

[10] K. Sugiyama, S. Tagawa, M. Toda, Methods for visual understanding of hierarchical system structures, IEEE Transactions on Systems, Man, and Cybernetics 11 (2) (1981) 109–125. doi:10.1109/TSMC.1981.4308636.

[11] P. Eades, N. C. Wormald, Edge crossings in drawings of bipartite graphs, Algorithmica 11 (1994) 379–403.

[12] D. Cheng Zarate, P. Le Bodic, T. Dwyer, G. Gange, P. Stuckey, Optimal Sankey diagrams via integer programming, 2018, pp. 135–139. doi:10.1109/PacificVis.2018.00025.

[13] K. M. Hall, An r-dimensional quadratic placement algorithm, Management Science 17 (3) (1970) 219–229. URL http://www.jstor.org/stable/2629091

[14] Y. Koren, Drawing graphs by Eigenvectors: Theory and practice, Computers Mathematics with Applications 49 (2005) 1867–1888. doi:10.1016/j.camwa.2004.08.015.

[15] J. N. Warfield, Crossing theory and hierarchy mapping, IEEE Transactions on Systems, Man, and Cybernetics 7 (7) (1977) 505–523.

[16] N. Liu, W. J. Stewart, Markov Chains and Spectral Clustering, Springer Berlin Heidelberg, Berlin, Heidelberg, 2011, pp. 87–98. doi:10.1007/978-3-642-25575-58. URL https://doi.org/10.1007/978-3-642-25575-5_8

[17] S. Mitchell, S. M. Consulting, I. Dunning, Pulp: A linear programming toolkit for python (2011).

[18] T. Herzog, World greenhouse gas emissions in 2005, World Resources Institute (2009).

[19] J. G. Caporaso, C. L. Lauber, E. K. Costello, et al., Moving pictures of the human microbiome, Genome Biology 12 (5) (2011) R50. doi: 10.1186/gb-2011-12-5-r50.

[20] L. Michielsen, M. J. T. Reinders, A. Mahfouz, Hierarchical progressive learning of cell identities in single-cell data, Nature Communications 12 (1) (2021) 2799. doi:10.1038/s41467-021-23196-8. URL https://doi.org/10.1038/s41467-021-23196-8

[21] C. Lee, C. Grasso, M. Sharlow, Multiple sequence alignment using partial order graphs, Bioinformatics 18 (3) (2002) 452–464. doi:10.1093/bioinformatics/18.3.452.

[22] A. Bahr, J. D. Thompson, J.-C. Thierry, O. Poch, Balibase (benchmark alignment database): enhancements for repeats, transmembrane sequences and circular permutations, Nucleic acids research 29 (1) (2001) 323–326.

